# Single-Dose Monocrotaline Pyrrole Injection as a Model of Pulmonary Endothelial Injury In Mice

**DOI:** 10.1101/305235

**Authors:** William Raoul, Anne Hulin, Guitanouch Saber, Catherine Boisnier, Saadia Eddahibi, Serge Adnot, Bernard Maitre

**Author notes:** Corresponding author: William Raoul, Unité INSERM U651 Faculté de Médecine de CRETEIL 8 rue du Général Sarrail, 94000 Creteil, FRANCE.

## Abstract

Monocrotaline (MCT) is a plant substance that induces severe pulmonary hypertension in several animals except for mice. The aim of our study was to state whether monocrotaline pyrrole (MCTp), the main monocrotaline metabolite, could induce significant injury in mouse lung when given intravenously. MCTp caused moderate pulmonary inflammation, remodelling of small distal vessels (percentage of muscularized arteries: 33,5 vs 20,6%, p≤0,0006) and a right ventricular dysfunction (RVSP 27,8mmHg vs 16,4mmHg, p≤0,0001; Fulton index 0,35 vs 0,26, p≤0,0007). These vascular effects were associated with a decrease in eNOS protein expression in lung tissues and resolved after 45 days. In conclusion, we developed a model of endothelial dysfunction and transient pulmonary hypertension in mice.

## Introduction

Pyrrolizidine alkaloids are toxins found in hundreds of plant species. Among them, monocrotaline (MCT) is found in the seeds and leaves of *Crotalaria spectabilis* (1, 2). When activated by the cytochrome P450 3A mono-oxygenase system in the liver to pyrrolic metabolites (3), MCT is toxic for both the liver and the lungs. Lung alterations induced by low-dose MCT include interstitial oedema, inflammation, and bleeding, with later addition of medial pulmonary artery hypertrophy, pulmonary hypertension, and right ventricular (RV) hypertrophy. In rats, MCT causes pulmonary hypertension with initial endothelial-cell injury (4). In contrast, mice given MCT continuously in drinking water for 6 weeks did not have marked pulmonary hypertension (as assessed by the ratio of RV weight over left ventricular [LV] plus septum weight [Fulton index], which reflects RV hypertrophy) (5); however, the highest dose (24 mg/kg/day) induced immune-mediated inflammation and mild endothelial dysfunction with decreases in angiotensin-converting enzyme, plasminogen activator activities, and prostacyclin and thromboxane production. This difference between the two species has been ascribed to MCT metabolism in the liver related to cytochrome P450 activity (6,7).

MCT pyrrole (MCTp or dehydromonocrotaline) is an active bifunctional alkylating agent that is thought to be the major hepatic MCT metabolite produced after MCT ingestion. When MCTp was given intravenously to rats (8), the toxic effects were closely similar to those of MCT. Consequently, we sought to develop a mouse model of MCTp injury after a single intravenous injection, based on various criteria including pulmonary lesion description, haemodynamic markers, inflammation markers, endothelial function marker (endothelial nitric oxide synthase expression).

Administration of MCTp in mice led to moderate pulmonary inflammation and marked remodelling of small pulmonary vessels with increases in RV systolic pressure and in the Fulton index. Endothelial function was also altered with transient dramatic decrease in eNOS protein expression. These results indicate that a single intravenous dose of MCTp induced transient pulmonary hypertension in mice associated with endothelial dysfunction.

## Material and Methods

### Study design

Male C57BL/6J mice (Charles River Laboratories) weighing 18 to 22 g were housed four per plastic cage in animal isolators (Charles River) under conditions of controlled temperature (25°C), humidity (15%) and light cycle (light:dark, 12:12). They were allowed food (Safe; Augy, France) and tap water *ad libitum.* All animal care and procedures were in accordance with institutional guidelines. A first series of mice was challenged with MCTp or vehicle to assess haemodynamic measurements and vascular remodelling in histology at different time points. Another series of mice was injected MCTp in order to characterize inflammation in broncho-alveolar lavage fluid and histopathology. A third set of experiments was performed to evaluate lung wet to dry weight ratio and to harvest lung samples for western blot analysis.

### MCTp synthesis and administration

Monocrotaline (Sigma-Aldrich, lot number # 07514DI, Sigma, purity 94%) was converted to MCTp by the method of Mattocks et al. (9). Identity and purity were assessed by high-pressure liquid chromatography and Ehrlich analysis as described by Molyneux and Roitman (10). MCTp was dissolved in N,N-dimethylformamide (DMF)/RPMI 1640 (Invitrogen, Cergy-Pontoise, France) just before use. Animals were placed in a plastic cylindrical chamber, anesthetized with isoflurane (Forene®, Abbott, Queenborough, UK), and given a single injection of MCTp (5 mg/kg) or the same volume of DMF/RPMI 1640 (control animals) in a tail vein (complete injection was ensured with slight aspiration of blood in syringe prior to product administration). Animals were killed 4 hours (H4), 1 day (D1), 5 days (D5), 10 days (D10), 15 days (D15), 21 days (D21), 45 days (D45), or 60 days (D60) after MCTp administration and used to evaluate markers for lung injury.

### Haemodynamic measurements and assessment of right ventricular hypertrophy

Mice were anesthetized and ventilated with room air at a tidal volume of 0.1 ml ketamine (7 mg/100 g) and xylazine (1 mg/100 g), at a rate of 90 breaths/min. After incision of the abdomen, a 26-gauge needle was inserted percutaneously into the right ventricle through the subxyphoid approach and right ventricular systolic pressure (RVSP) was recorded immediately using a Gould pressure transducer (Gould P23 ID, Gould Electronics, France) connected to pressure modules and a Gould TA 550 recorder. Monitoring of RVSP profile was also done with an oscilloscope (Hameg, Francfort, Germany). After an intraperitoneal injection of sodium pentobarbital (40 mg/kg) and exsanguination, the thorax was opened and the lungs and heart were removed. The RV was dissected from the LV and septum, and these samples were weighed; Fulton's index was computed as the ratio of RV weight over LV plus septum weight and was used to assess RV hypertrophy.

### Pulmonary vascular remodelling

In each mouse, a total of 30 intra-acinar vessels (25 to 44 μm) accompanying either alveolar ducts or alveoli were examined by an anatomopathologist blinded to treatment allocation. Each vessel was categorized as non-muscular (either no evidence of vessel wall muscularisation or smooth muscle cells (SMCs) in less than half the vessel circumference) or muscular (SMCs in more than half the vessel circumference). Muscularization was defined as the presence of typical SMCs stained red with phloxin, exhibiting an elongated shape and square-ended nucleus, and bound by two orcein-stained elastic laminae. The percentage of pulmonary vessels in each muscularisation category was determined by dividing the number of vessels in that category by the total number counted in the same experimental group.

### Evaluation of inflammation

#### Bronchoalveolar lavage (BAL) and cytospin

After lung harvesting, a 0.6-mm tube was inserted through a small incision in the trachea. For BAL, 2 ml of 9 %o NaCl (1x 1 ml and 2x 0.5 ml) was instilled in the tube then gently aspirated. The samples were then centrifuged at 300 g for 7 minutes at 4 °C, and the cell-free BAL fluid was stored at −70°C.

Pellets were then resuspended with the remaining supernatants, and cell numbers were assessed with a haemocytometer. Cytospin preparations were obtained by placing 0.2-ml BAL aliquots in each funnel of a Cytospin III (Shandon, Cergy-Pontoise, France). The cells were fixed and stained with Diff-Quick (Dade Behring, La Défense, France), and a differential leukocyte count was performed.

#### Histology

Bronchoalveolar lavage (BAL) fluid was collected and the lungs were fixed by intratracheal infusion of 4% paraformaldehyde in phosphate-buffered saline (PBS, Invitrogen, Cergy-Pontoise, France) at a pressure of 23 cm H2O. After 24 h, the tissues were dehydrated and embedded in paraffin then cut into 5-μm slices, which were stained with hematoxylin-phloxin-saffron and orcein-picroindigo-carmine for histological examination. The inflammatory response was assessed using a previously described empiric semi-quantitative scale (11) based on inflammatory-cell type and location (alveoli, bronchi, or blood vessels), oedema, and bleeding. The extent of inflammation was scored on a 0-4 scale as follows: 0, none; 1, small scattered areas; 2, <10%; 3, 10-50%; and 4, >50% of section area.

#### Quantification of lung wet/dry weight ratio

Another series of lungs were excised en bloc and dissected away from the heart and thymus. These lungs were immediately weighed (wet weight) then placed in a desiccating oven at 65 °C for 48 h and weighed again (dry weight). The wet/dry weight ratio was used to quantitate the lung water content (12).

### Endothelial nitric oxide synthase (eNOS) expression by western blot analysis

Removed lungs were quickly frozen in liquid nitrogen. After thawing at 0 °C, the lung tissues were sonicated in 0.1 mM PBS containing anti-proteases (1 μM leupeptin and 1 μM pepstatin) and CHAPS (20 mmol/L). The homogenate was centrifuged at 3000 g for 10 min at 4 °C. Direct protein quantitation was done by the Bradford method with IgG standardization purchased from Bio-Rad (Ivry-sur-Seine, France). Detection of eNOS was by western blot as described by Pascaud *et al.* (13). Briefly, after polyacrylamide gel electrophoresis of the samples (200 μg each) and transfer to a PVDF membrane by electroblotting (12h, 4 °C), the membrane was blocked with TTBS (0.15 M NaCl, 1 mM Tris-HCl, pH=8, 0.05% Tween 20, and 5% bovine serum albumin [BSA]) for 1 h at room temperature, and eNOS protein was detected using a mouse monoclonal anti-eNOS primary antibody reactive with mouse, rat or human eNOS (Transduction Laboratories, Lexington, USA), diluted 1:1000. Then, specific protein was detected using a horseradish peroxidase-conjugated secondary antibody (Calbiochem, San Diego, USA), diluted 1:10,000 and ECL reagents (Amersham, Orsay, France). Quantitation of eNOS immunoreactivity was achieved using a semi-automated image analysis device (GeneTools software, Syngene, Cambridge, UK). Results are reported in arbitrary units.

### Statistical analysis

All results are reported as means ± standard error of the mean (SEM). MCTp effects at various time points were compared using Kruskall Wallis test for analysis of variance, followed by a Mann Whitney test to compare each time point with the control. Statistical significance was defined as *P*<0.05. Statistical analysis was performed using STATVIEW 5.0 software (Abacus Concepts, Inc, Berkeley, CA).

## Results

Based on rat MCTp data, we decided to use three different doses in mice: 5 mg/kg, 10 mg/kg and 20 mg/kg. These different doses induce a same increase of RVSP and Fulton index on day 15 (data not shown). We then chose the 5 mg/kg dose for all experiments described above.

### Haemodynamic parameters and assessment of right ventricular hypertrophy

RVSP (figure 1A) and Fulton's index (figure 1B) were similar in the control group (vehicle DMF/RPMI 1640) and in the MCTp-treated group until D5. Animals given a single injection exhibited a significant increase in RVSP and in the Fulton index on D10 and D15, as compared with the control group. Haemodynamic parameters are equivalent between D15 and D21. In the longer term, no differences were found between control and MCTp groups on D45 and D60.

**Figure 1.**
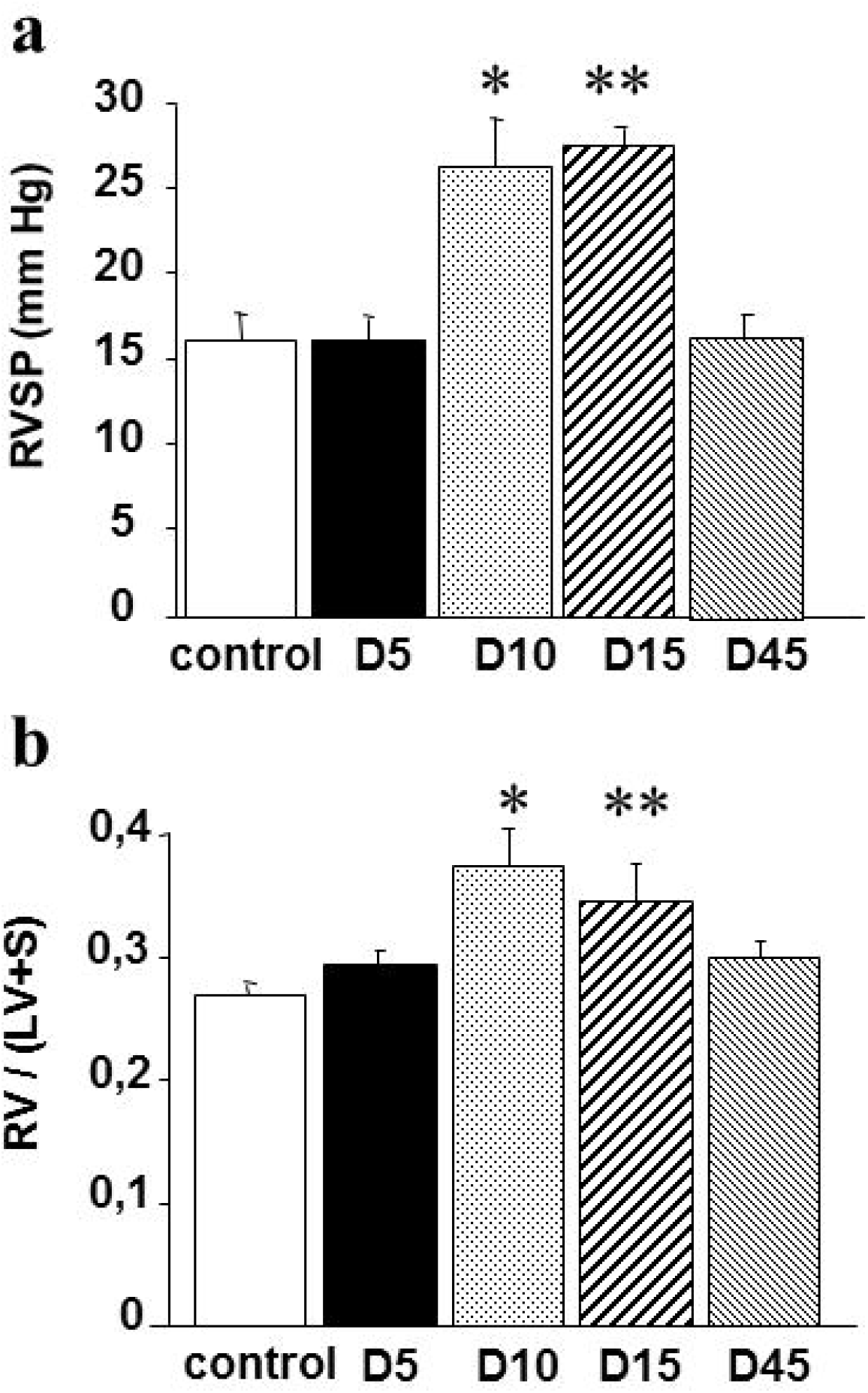
a. Right ventricular systolic pressure (RVSP) in mice at various times after administration of a single 5-mg/kg dose of monocrotaline pyrrole (MCTp) (black bars) and in control mice not given MCTp (open bar). \**p*≥0.004, \*\**p*≥0.0006 for comparisons with values in control mice. Values are means±SEM for n=5 animals in each group. b. Ratio of right ventricle to left ventricle + septum weight [RV/(LV+S), Fulton index] in mice at various times after administration of a single dose of 5 mg/kg monocrotaline pyrrole (MCTp) (black bars) and in control mice not given MCTp (open bar). \**p*≥0.0001, \*\**p*≥0.01 for comparisons with values in control mice. Values are means±SEM for n=6-10 animals in each group.

### Structural remodelling of distal pulmonary vessels

In MCTp-treated mice as compared to controls, muscularisation of distal pulmonary vessels was similar until D10, markedly increased on D15 and D21, and similar on D45 and D60 (figure 2).

**Figure 2.**
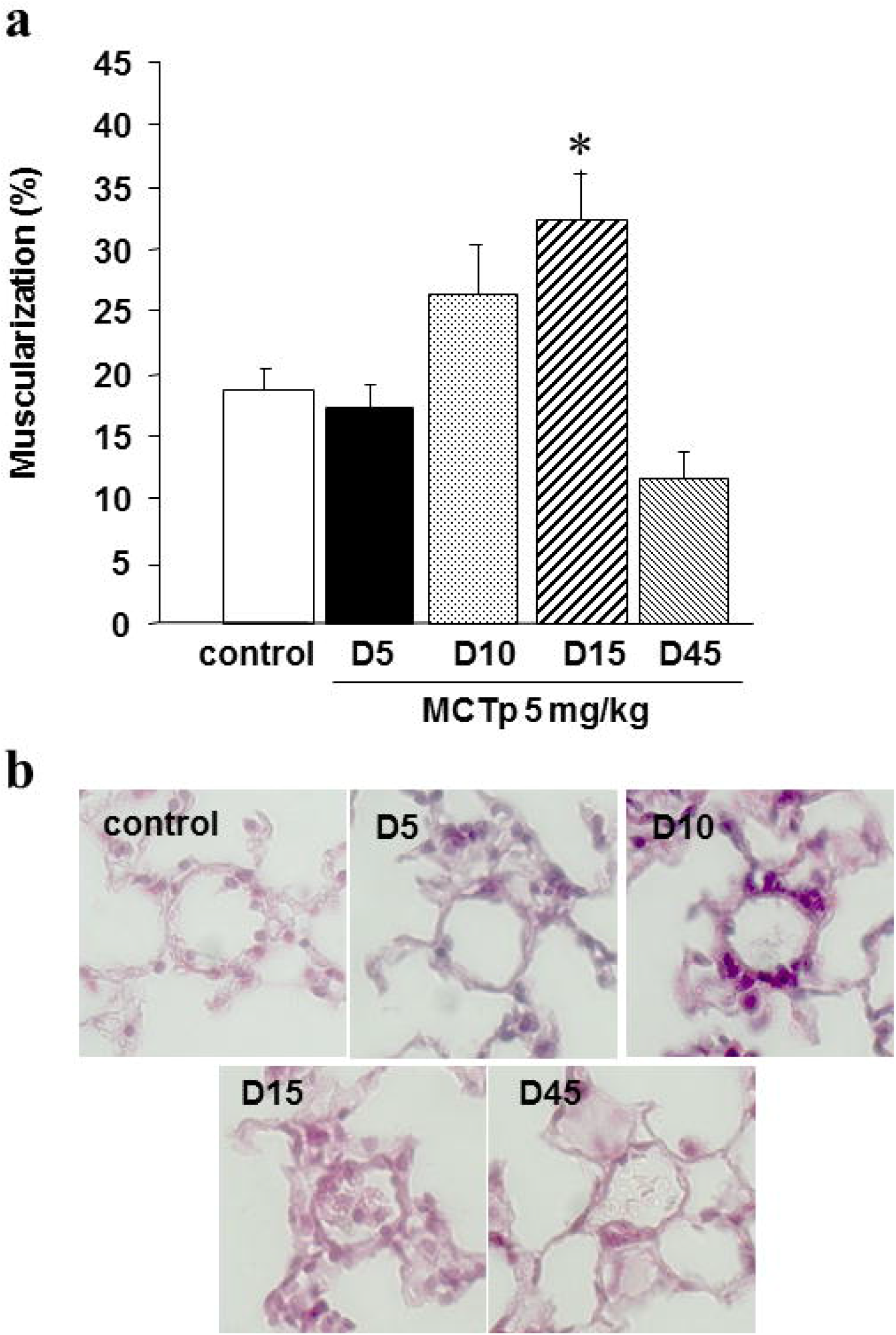
a. Percentage of muscularised intra-acinar vessels in lungs from mice at different times after administration of a single dose of 5 mg/kg monocrotaline pyrrole (MCTp) (black bars) and in control mice not given MCTp (open bar). \**p*≥0.0007 for comparisons with values in control mice. Values are means±SEM for n=5 animals in each group. b. Representative panels comparing muscularised distal vessels and non-muscularised distal vessels. Lung morphology in control mice (C), 5 days (D5), 10 days (D10), 15 days (D15) and 45 days (D45) after a single injection of 5 mg/kg MCTp. Sections were stained with hematoxylin-phloxin-saffron. Original magnification x500.

### Evaluation of lung inflammation

#### Histologic results

At H4 and D1, marked macrophage recruitment was noted near proximal arteries and around bronchi (panels a and b,figure 3), and inflammatory infiltrates with interstitial macrophages were seen in the alveolar walls. On D5, inflammatory cells were still present in the alveolar spaces but inflammation mainly affected the proximal arteries, some of which exhibited elastic-fibre alterations (panel c, figure 3). On D10 and D15, inflammation was less marked and progressive vascular remodelling was seen as medial thickening of the proximal arteries and muscularisation of the distal pulmonary vessels (panels d and e, figure 3). On D60, lung structure was normal with few macrophages, no medial thickening in proximal arteries, and reduced muscularisation of distal vessels.

**Figure 3.**
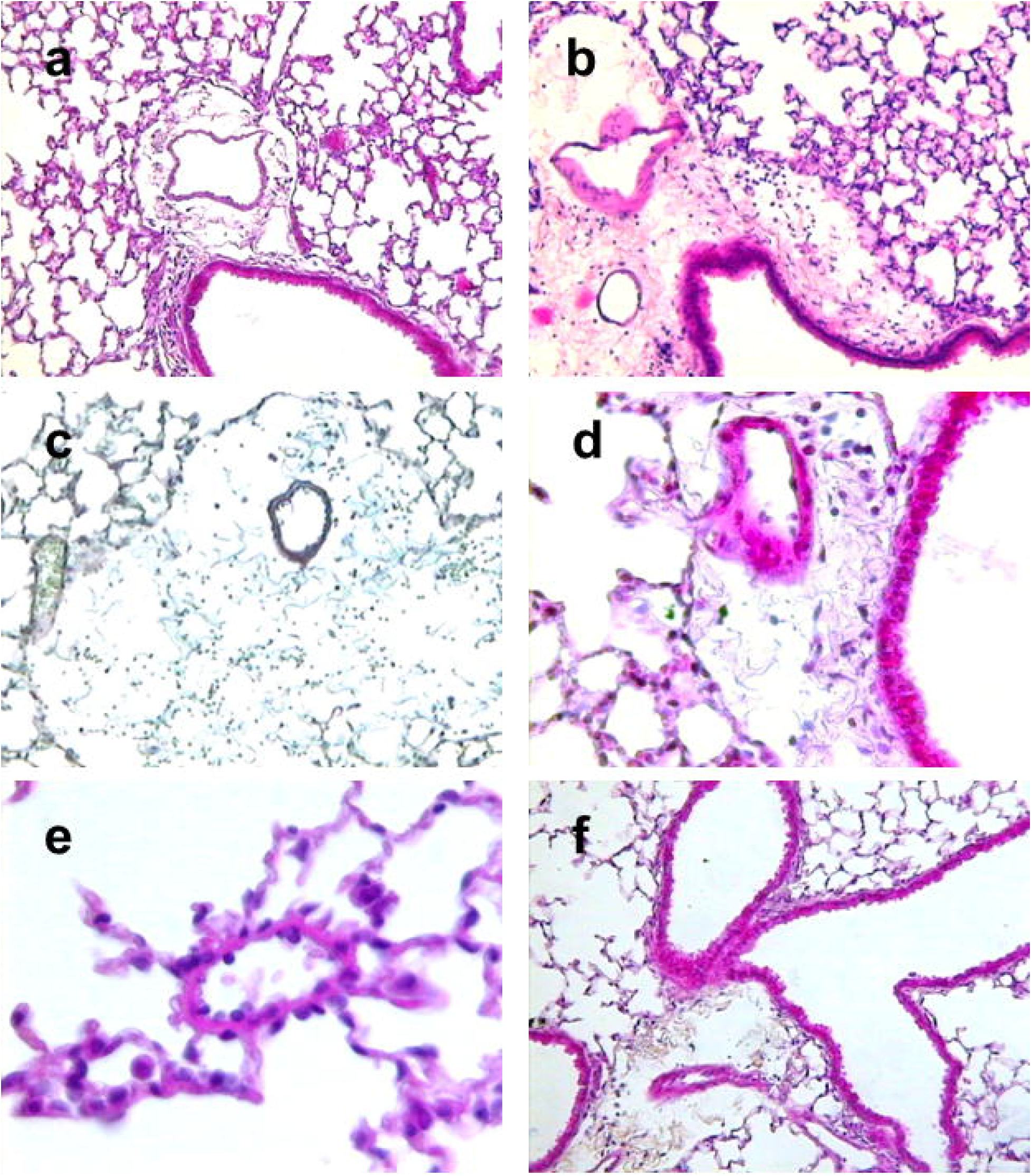
Lung histology in mice given a single injection of 5 mg/kg MCTp. Sections were stained with hematoxylin-phloxin-saffron excepted panel c with orceine-picroindigo-carmine. (Original magnification a,b,c,f: x125; d: x250; e: x500) Panels a and b: representative slides showing inflammatory cell recruitment at 4 hours and 24 hours after MCTp administration, respectively. This inflammation is localized mainly in peribronchial and alveolar area. Panel c: representative slide showing persistent inflammation and injury of elastic fibres (altered blue staining) on day 5 after MCTp administration. Panels d and e: representative slides showing vascular remodelling of bronchial arteries and of distal pulmonary vessels with typical red phloxin staining of smooth muscle cells, respectively, with a decrease in inflammatory-cell infiltrates on day 15 after MCTp administration. Panel f: representative slide showing normal pulmonary structure on day 45 with very few inflammatory cells and no increase in media thickness of bronchial arteries.

Table 1 reports the extent of inflammation. Diffuse airway inflammation and macrophages in proximal arteries were noted from H4 to D5. The inflammation abated gradually, and parameters were back to baseline values on D45 and D60. In our model, the livers were normal by gross examination or light microscopy.

**Table 1. Inflammation in lung histopathology and in broncho-alveolar lavage fluid.**

|  | Control | H4 | D1 | D5 | D10 | D15 |
| --- | --- | --- | --- | --- | --- | --- |
| **Lung Histopathology** |  |  |  |  |  |  |
| Inflammatory cells | mφ | mφ | mφ | mφ | mφ | mφ |
| Inflammation extent<br>(0 to 4) | 0 | 3 | 2 | 2 | 2 | 1 |
| Main location | - | prox | prox | prox/dist | prox/dist | prox |
| **BAL fluid** |  |  |  |  |  |  |
| Inflammatory cells | mφ | mφ | mφ | mφ | mφ | mφ |
| Total cell count<br>(cells x10<sup>5</sup>) | 1±0,3 | 1,8±0,4 | 1,3±0,6 | 1,8±0,8 | 1,4±0,4 | 1,1±0,3 |

Definition of abbreviations: H, hour; D, Day; mφ, macrophages; prox, proximal arteries; dist, distal vessels and alveolar cells; BAL, broncho-alveolar lavage. The extent of inflammation was scored as follows: 0, none; 1, small scattered areas; 2, <10%; 3, 10-50%; 4, >50% of section area. n = 5 animals / group

#### BAL fluid

There was no significant increase in alveolar macrophage numbers at H4 after MCTp administration as compared to the control group. The number of macrophages in BAL fluid only tended to increase then decrease steadily from H4 onward, reaching control values by D15 (table 1). However, no increase in polymorphonuclear cells was noted.

#### Oedema

Oedema evaluated by wet-to-dry weight ratio showed no difference at any time point between the control group and the MCTp-treated groups (control: 3.9±0.1; and MCTp: 4.1±0.2 on D5, 3.6±0.1 on D10, and 3.3±0.1 on D15).

### Effect of MCTp on eNOS expression

To evaluate the consequences of MCTp administration on endothelial markers, lung eNOS level was estimated by immunoblotting at various time points. As shown in figure 4, eNOS level was significantly lower on D15 after MCTp injection than in the control group. Levels were identical on D15 and D21 and were restored on D45.

**Figure 4.**
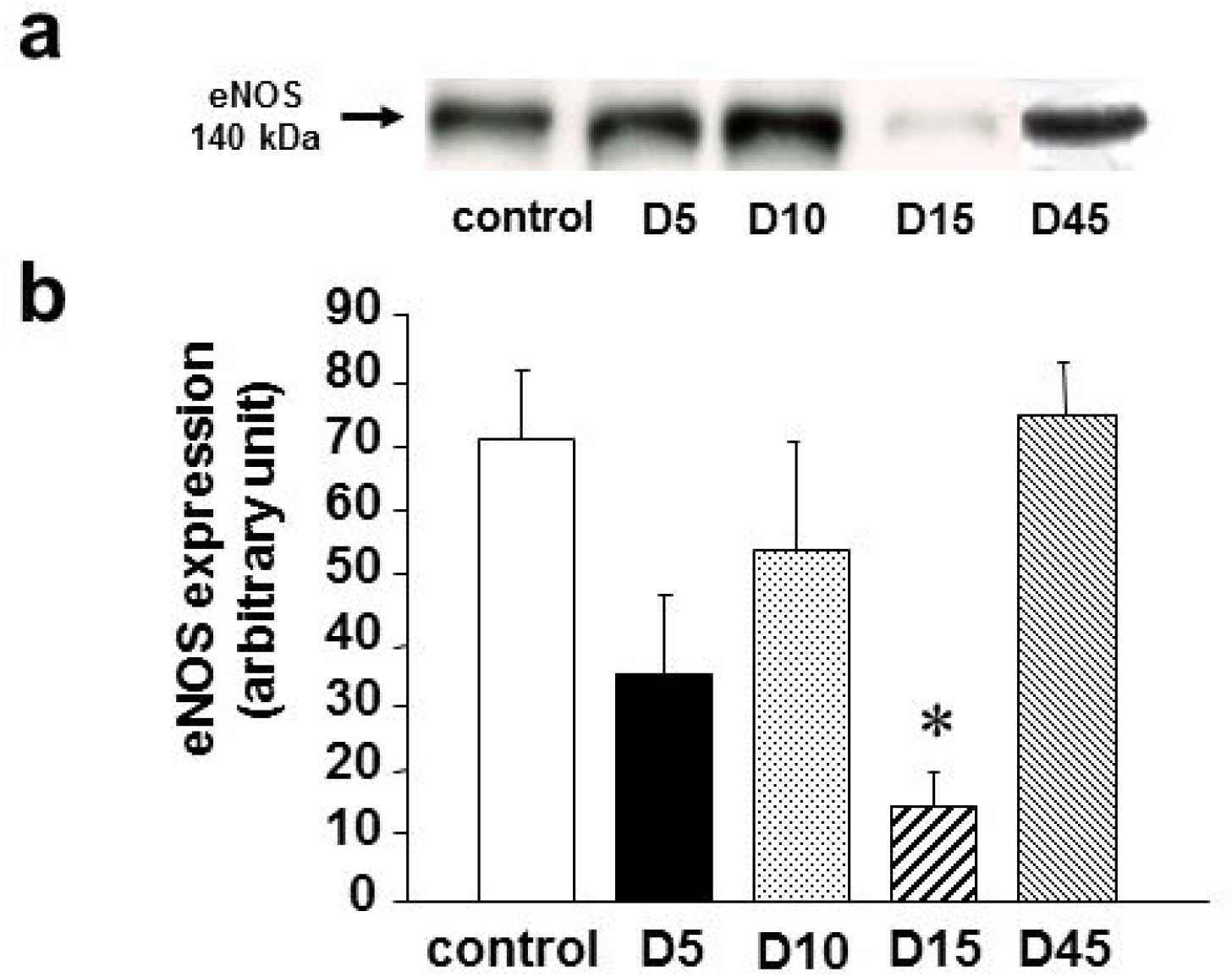
Western blot analysis of the effects of 5 mg/kg MCTp on endothelial nitric oxide synthase (eNOS) protein expression by mouse lung tissues. (a) Representative western blots at various times following MCTp administration compared with control mice (control) not given MCTp. (b) Histograms of eNOS quantities in lung tissues from control mice (open bar) and MCTp- treated mice (black bars). \**p*≥0.01 for comparison with values in control mice. Values are means±SEM for n=4 animals in each group.

## Discussion

The histologic and haemodynamic findings from our studies show clearly that transient pulmonary hypertension developed in the MCTp-treated mice. Thus, on D15, increases were noted in the Fulton index (indicating RV hypertrophy) and in RVSP, and histology showed muscularisation of distal pulmonary vessels; in addition, endothelial cell activity was significantly altered by the initial injury, with major decrease in eNOS protein expression by western blot analysis. The abnormalities were greatest on D15 and resolved subsequently.

Pulmonary hypertension was present from D10 onward in our mouse model, as compared to D8 in rats, indicating that the time-course of pulmonary hypertension development was roughly similar in the two species. Schultze *et al.* (14) showed an increase in Fulton index from 0.30 to 0.48 (day 14) in rats treated with MCTp. Our percentage of increase of Fulton index was closed (0.26 to 0.38) and it has been noticed previously that mouse models of pulmonary hypertension are generally less severe than rat models (15).

Since there is no equivalent model of lung injury in mice, we will compare our data to those from models of pulmonary artery remodelling, such as hypoxia-induced pulmonary hypertension. MCTp induced a similar pattern of remodelling. Muscularisation of distal pulmonary vessels was slightly less marked in our model as compared to hypoxia-induced pulmonary hypertension. In a model of chronic hypoxia induced-pulmonary hypertension, Ozaki *et al.* (16) with the same RVSP evaluation (right ventricle puncture) and the same strain (C57BL/6) found RVSP values from 22 mm Hg in normoxia to 34 mm Hg in hypoxia, Fulton index evolved from 0.29 to 0.42. These findings corroborated our data showing almost the same scale of values. Furthermore, there was little or no inflammation in hypoxia-induced pulmonary hypertension, and pulmonary artery pressure returned to normal when the animals were switched back to normoxia.

Several studies focused on MCTp-induced injury in rats and showed delayed and gradual development of inflammatory events. In a study by Roth and Reindel (17) that relied mainly on electron microscopy, vascular leakage and pulmonary oedema were visible on day 3 post-injection; small artery remodelling and pulmonary inflammation on day 5; and platelet sequestration, large artery remodelling, and pulmonary hypertension on day 8, on day 14 the observations were stopped. Schultze *et al.* (18) found significant increase in neutrophil recruitment at D5 after MCTp administration to rats, as well as oedema. Our results showed no evidence of inflammation in BAL fluids and oedema assessment was not a prominent feature but histological studies showed clearly interstitial macrophages recruitment. We did not assess inflammation markers after peak injury on D15 because inflammation parameters returned to baseline values at this time. Morerover, we showed gradual development of transient pulmonary hypertension without massive mononuclear infiltration into perivascular lesions. Differences in susceptibility to inflammation between rats and mice studies may be ascribable to differences in metabolism and enzyme pools. It will be interesting to test the effects of anti-inflammatory strategies in this mouse model since several articles report a beneficial effect of these strategies in the rat monocrotaline model (19).

To gain some insight into endothelial function, we decided to assess the protein expression of eNOS in this model. NO bioactivity and bio disponibility are critical for development of pulmonary hypertension (20) and eNOS is expressed mainly in endothelial cells and is a endothelial marker. eNOS protein expression was dramatically decreased in our model but was considerably increased in chronic hypoxia-induced pulmonary hypertension in mice (13), suggesting a key role for endothelial dysfunction in MCTp-induced pulmonary hypertension.

In conclusion, we developed a model of monocrotaline pyrrole-induced endothelial dysfunction and transient pulmonary hypertension in mice. Early moderate inflammation developed in the lung vessels and alveolar walls, followed by progressive remodelling of small distal arteries with development of marked pulmonary hypertension. Moreover, endothelial function was clearly altered. This model will be useful to test potential treatments for endothelial dysfunction and/or transient pulmonary hypertension in mice over a long period (about 2-3 weeks before spontaneous resolution of the abnormalities). Furthermore, this model may prove useful for transgenic approaches and progenitor cells research.

